# Effectiveness and relationship between biased and unbiased measures of dopamine release and clearance

**DOI:** 10.1101/2021.12.20.473490

**Authors:** Anna C. Everett, Ben E. Graul, J. Kayden Robinson, Daniel B. Watts, Rodrigo A. España, Cody A. Siciliano, Jordan T. Yorgason

## Abstract

Fast-scan cyclic voltammetry (FSCV) is an effective tool for measuring dopamine (DA) release and clearance throughout the brain, including the ventral and dorsal striatum. Striatal DA terminals are abundant with signals heavily regulated by release machinery and the dopamine transporter (DAT). Peak height is a common method for measuring release but can be affected by changes in clearance. The Michaelis-Menten model has been a standard in measuring DA clearance, but requires experimenter fitted modeling subject to experimenter bias. The current study presents the use of the first derivative (velocity) of evoked DA signals as an alternative approach for measuring dopamine release and clearance and can be used to distinguish the two measures. Maximal upwards velocity predicts reductions in DA peak height due to D_2_ and GABAB receptor stimulation and by alterations in calcium concentrations. The Michaelis-Menten maximal velocity (V_max_) measure, an approximation for DAT numbers, predicted maximal downward velocity in slices and *in vivo*. Dopamine peak height and upward velocity were similar between wildtype C57 (WT) and DAT knock out (DATKO) mice. In contrast, downward velocity was considerably reduced and exponential decay (tau) was increased in DATKO mice, supporting use of both measures for changes in DAT activity. In slices, the competitive DAT inhibitors cocaine, PTT and WF23 increased peak height and upward velocity differentially across increasing concentrations, with PTT and cocaine reducing these measures at high concentrations. Downward velocity and tau values decreased and increased respectively across concentrations, with greater potency and efficacy observed with WF23 and PTT. In vivo recordings demonstrated similar effects of WF23 and PTT on measures of release and clearance. Tau was a more sensitive measure at low concentrations, supporting its use as a surrogate for the Michaelis-Menten measure of apparent affinity (Km). Together, these results inform on the use of these measures for DA release and clearance.

## Introduction

Fast-scan cyclic voltammetry (FSCV) is a widely used electrochemical detection technique that can be highly selective for specific analytes. This technique is perhaps most frequently used for detection of monoamines (such as dopamine) and indolamines (For review see (Ferris *et al*. 2013)), but is also increasingly used for detecting other neurotransmitters such as hydrogen peroxide and purines (Li & Ross 2020) (Spanos *et al*. 2013). Fast-scan cyclic voltammetry is powerful because of its high temporal kinetics and chemical specificity, making it an ideal tool for studying the mechanisms underlying rapid neurotransmitter release and clearance. For the past 30 years, the Michaelis-Menten model has been used alongside voltammetry techniques to study the kinetics of the dopamine transporter (DAT) (Wightman 1988) (Wightman & Zimmerman 1990) (Yorgason *et al*. 2011a) (Wu *et al*. 2001). In general, the Michaelis-Menten equation is used to measure first order enzymatic kinetics, solute binding efficiency, and enzyme concentrations (Michaelis *et al*. 2011). The maximal rate of dopamine uptake is referred to as Vmax while the DAT’s apparent binding affinity for dopamine is known as the Michaelis-Menten constant K_m_. The voltammetry modified version of this model was described previously (Wightman & Zimmerman 1990) (Yorgason *et al*. 2011a) and includes the typical V_max_ and K _m_ variables, in addition to variables describing the release concentration and a thickness layer. Release concentration is used to describe the amount of DA detected, and the thickness layer variable is a deconvolution factor, which is often used as a correction coefficient to better match biological signals.

While this model has excellent predictability for measuring differences in dopamine transporter function, calculating baseline measures of V_max_ are highly subject to experimenter bias. Thus, the present work examined alternative methods for studying DA release and clearance kinetics involving the first (velocity) and second (acceleration) derivative, which are easily replicable measures of slope for a signal. Velocity and acceleration values were examined in conditions that are known to influence DA release and clearance. These measures were also compared to Michaelis-Menton model-based measures of V_max_ from slice and *in vivo* preparations. Unlike the Michaelis-Menten model, these analytical measures are not dependent on enzymatic activity, making them ideal for describing clearance in animal models that do not contain transporters. Thus, the present work describes this use of easily measurable values of slope that have reduced analytical bias and may inform on applications for studying diffusion and other non-DAT-mediated DA clearance.

## Materials & Methods

### Animals

Male and Female (Jackson Laboratory, Sacramento, CA) WT and DATKO (~25-35 g) mice on C57Bl/6 background were given ad libitum access to food and water and maintained on a 12:12-h light/dark cycle. All protocols and animal care procedures were in accordance with the National Institutes of Health Guide for the Care and Use of Laboratory Animals and approved by Brigham Young University Institutional Animal Care and Use Committee, Oregon Health and Science, Drexel University College of Medicine, and Wake Forest University Health Sciences.

### Brain Slice Preparation and Drug Application

Isoflurane (Patterson Veterinary, Devens, MA) anesthetized mice were sacrificed by decapitation and brains were rapidly removed, sectioned into 220-400 μm-thick coronal striatal slices (Leica VT1000S, Vashaw Scientific, Norcross, GA), and incubated for 60 minutes at 34 °C in preoxygenated (95% O2/5% CO2) artificial cerebral spinal fluid (aCSF) consisting of (in mM): NaCl (126), KCl (2.5), NaH2PO4 (1.2), MgCl2 (1.2), NaHCO3 (25), D-glucose (11), L-ascorbic acid (0.4), with pH adjusted to ~7.4. Cutting solution also contained either MK801 (10 μM; (5S,10R)-(+)-5-methyl-10,11 -dihydro-5H-dibenzo[a,d]cyclohepten-5,10-imine; Abcam, Cambridge, UK) or kynurenic acid (2 mM) for blockade of ionotropic glutamate receptors. At the end of the incubation period, the tissue was transferred to aCSF (34 °C) without glutamate receptor blockers. The following concentrations of drugs were bath applied for slice voltammetry experiments where specified: Baclofen (30 μM), CGP55845 (200 nM), quinpirole (10 μM), sulpiride (600 nM; Sigma), 2β-propanoyl-3β-(2-naphthyl)-tropane (WF-23) (10 nM – 3 μM; Huw M. L. Davies, Emory University, Atlanta DA), 2β-propanoyl-3β -(4-tolyl)-tropane (PTT) (100 nM – 30 μM; Huw M. L. Davies), cocaine (300 nM - 30 μM; NIDA, Rockville, MD USA).

### Ex Vivo Voltammetry Recordings

Slices were transferred to the recording chamber, and perfused with aCSF (34 °C) at a rate of ~1.8 ml/min. Fast scan cyclic voltammetry recordings were performed and analyzed using Demon Voltammetry and Analysis software (Yorgason *et al*. 2011a). Carbon fiber electrodes used in voltammetry experiments were made in-house. The carbon fiber (7 μm diameter, Thornel T-650, Cytec, Woodland Park, NJ) was aspirated into a borosilicate glass capillary tube (TW150, World Precision Instruments, Sarasota, FL). Electrodes were then pulled using a pipette puller and cut so that 100-150 μm of carbon fiber protruded from the tip of the glass. The electrode potential was linearly scanned as a triangular waveform from −0.4 to 1.2 V and back to −0.4 V (Ag vs AgCl) with a scan rate of 400 V/sec, repeated every 100 ms. Carbon fiber electrodes were positioned ~250 μm below the slice surface. For baclofen and quinpirole experiments, dopamine release was evoked through electrical stimulation (1 pulse/min) via a glass micropipette (30 μA, 0.5 ms), repeated every 2 minutes. For calcium and DATKO experiments, dopamine release was evoked from a bipolar stimulating electrode (Plastics One, Roanoke, VA) and used stimulations at 1 pulse, 5 pulses at 20 Hz, or 24 pulses at 60 Hz (300 μA, 4 msec) applied to the tissue every 5 minutes. Multiple pulse stimulation parameters were selected to model the different in vivo firing patterns of dopamine neurons (Daberkow *et al*. 2013) (Zhang *et al*. 2009). For baseline collections, stimulations were applied every 2 or 5 minutes until a stable baseline was established (3 collections within 10% variability). Next, either drug or stimulation conditions were changed as noted in the results.

### In Vivo Voltammetry Recordings

On the day of testing, mice were anesthetized with urethane (1.5 g/kg, i.p.; Sigma-Aldrich, St. Louis, MO, USA) and implanted with an intravenous (i.v.) catheter as previously described in (Yorgason *et al*. 2011b). All drugs for *in vivo* experiments were dissolved in 0.9% saline. Mice were subsequently placed in a stereotaxic apparatus while a stimulating electrode was lowered into the ventral tegmental area (VTA) (−3.0 A/P, +1.0 M/L, −4.5 – 5.0 D/V), a carbon fiber electrode was lowered into the dorsal caudate (+1.3 A/P, +1.3 M/L, −3.0 D/V) and a reference electrode was implanted in the contralateral cortex (+1.5 A/P, −1.5 M/L, −2.0 D/V) (Shaw *et al*. 2017). DA release was elicited via electrical stimulation of the VTA using 0.4 – 1 sec, 60 Hz monophasic (24p, 4 ms; ~400 μA) stimulation trains. The length of stimulation was varied to ensure sufficient DA release across experiments. Baseline DA response parameters were collected in 5 minute intervals for a minimum of 30 minutes prior to drug injection. Once a stable baseline of three consecutive collections was obtained (defined as DA peak height within 15%), mice received a single 2 second, ~200 μL i.v. injection of WF-23 (5.0 mg/kg n = 6), or PTT (5.0 mg/kg n = 6). DA response parameters were acquired at 30 and 60 seconds post injection and every 5 minutes thereafter until a peak effect was established.

### Data Analysis and Statistics

The magnitude of electrically-evoked DA release (peak height) and transporter-mediated uptake were monitored and DA overflow curves were fitted to an exponential decay model and expressed as tau (time to ~33% of peak height; units in seconds), then modeled using the Michaelis-Menten fit or using the fastest upward or downward velocity calculated from the first derivative of the release trace using Demon Voltammetry and Analysis software (Yorgason *et al*. 2011a) written in LabVIEW (National Instruments, Austin, TX).

Carbon fiber electrodes were initially calibrated using low (1 μM) and high (10 μM) DA concentrations, which was compared back to background current amplitude for establishing a model based on linear regression for calibration values (Supplementary methods and Supplementary Figure 1). The resultant linear regression model was applied to electrodes for determining the relative calibration constant. Statistics were performed using Prism 5 (GraphPad, San Diego, CA) and NCSS 8 (NCSS LLC, Kaysville UT). Statistical significance was determined for groups of 2 variables using a two-tailed student t-test. Experiments with more than 2 groups but only one factor were tested for significance using a one-way analysis of variance (ANOVA). Data from multiple time points within the same experiment were analyzed with a repeated measures ANOVA (RM-ANOVA). For experiments that examined multiple factors and interactions, two-way ANOVA or RM-ANOVAs were used. Tukey’s HSD and Bonferroni correction methods were used for post-ANOVA analysis.

## Results

### Upward velocity and peak height as measures of changes in dopamine release

In order to establish alternate measures of dopamine release, the maximal upwards velocity was examined under conditions that alter release probability. The release curve and first derivative of an evoked dopamine signal (1 pulse) are shown (**Figure 1A**). Dopamine release was examined before and after activation of two inhibitory G_i_-protein coupled receptors (GPCRs), GABA_B_, and D_2_ inhibitory receptors (**Figure 1B-C**). A maximal concentration of the GABA_B_ agonist baclofen (30 μM) reduced NAc dopamine peak height by 30.1%, which reversed upon antagonist application (CGP55845; 200 nM). The D2 agonist quinpirole (1 μM) reduced dopamine release to a much greater extent (92.7%), which similarly reversed with D2 antagonist application (sulpiride; 600 nM). Peak height and velocity up strongly correlated in these experiments (**Figure 1C**; n = 4, r^2^ = 0.9999, slope = 5.006 ± 0.0075).

**Figure 1.**
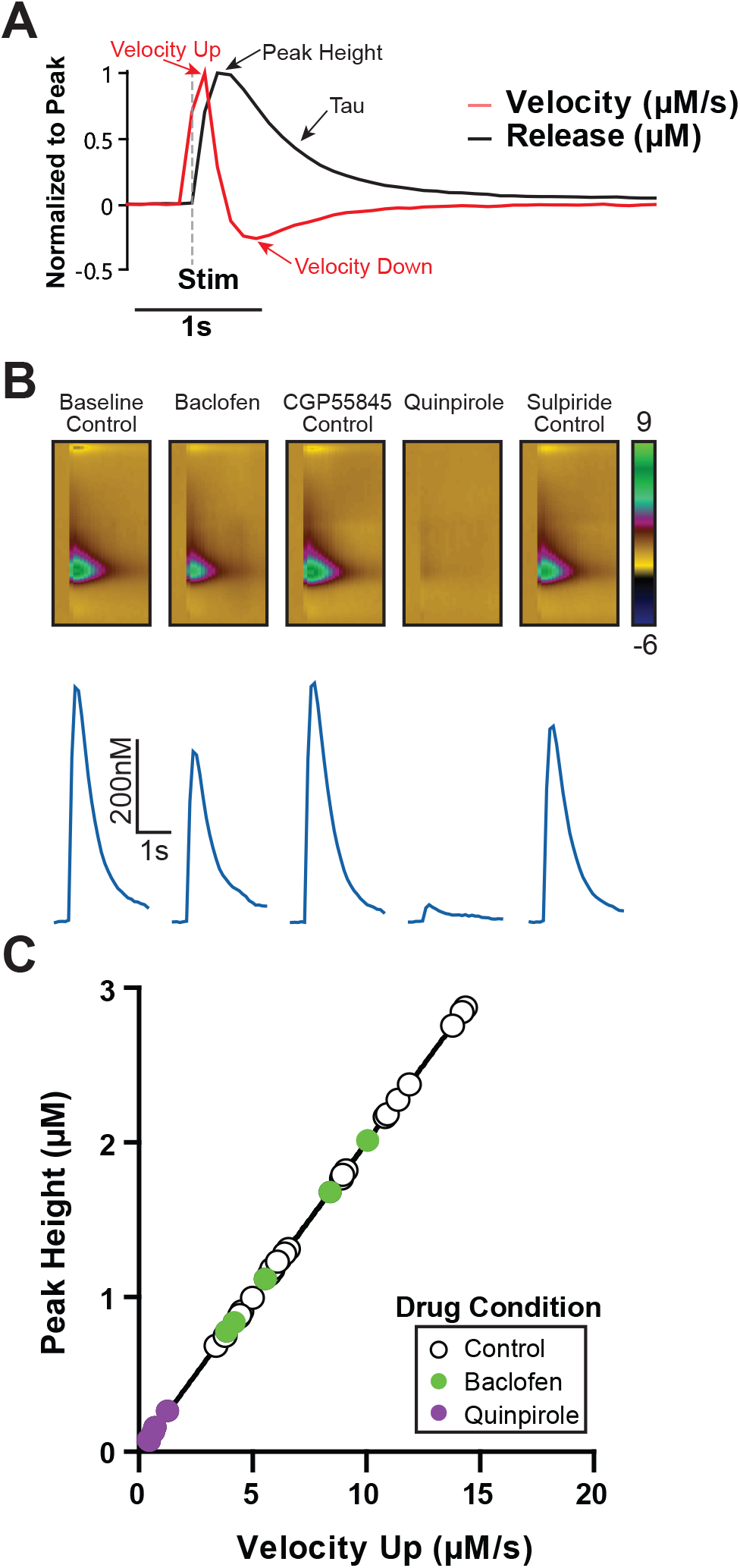
Peak height and upward velocity are related measures of dopamine release and are reduced with D_2_ and GABA_B_ agonists. A) An example dopamine trace is shown under baseline conditions (black), alongside its corresponding first derivative (red; velocity), normalized by peak to highlight how these measures correspond to each other. Time of stimulation (1p, 30 μA) is denoted (gray dashed line). B) Dopamine release is reduced by GABA_B_ agonist baclofen (30 μM), which is reversed by the GABA_B_ antagonist CGP55845 (200 nM), which acts as a control for baclofen effects. The D_2_ agonist quinpirole (1 μM) robustly inhibits dopamine release, which is reversed by the D_2_ antagonist sulpiride (600 nM), which was also treated as a control condition. C) Dopamine release peak height correlates with maximal upward velocity, which is decreased from control conditions during GABA_B_ and D_2_ stimulation.

Electrically evoked dopamine release is calcium dependent (Yorgason *et al*. 2014) (Ford *et al*. 2010) and dopamine release at physiological calcium (1.2 mM) levels is ~50% of release observed from maximal (2.4-4.8 mM) calcium concentrations (Karkhanis *et al*. 2019). Therefore, the effects of calcium (1.2-4.8 mM) on evoked dopamine release (20Hz, 5p) were compared between peak height and upward velocity measures (**Figure 2**). Experiments with higher calcium (2.4-4.8 mM) concentrations exhibited ~232% greater dopamine release (200% for 2.4 mM, 263% for 4.8 mM) than physiological (1.2 mM) calcium levels (**Figure 2A-B**; One-way ANOVA, n_1.2_ = 6 slices, n_2.4_ = 10 slices, n_4.8_ = 4 slices, *F*_2,17_ = 13.65, *p* < 0.0003). A Tukey’s HSD posttest revealed significantly increased dopamine release differences between calcium concentrations (1.2 to 2.4 mM: *p* < 0.01, q = 6.238; 1.2 to 4.8 mM: *p* < 0.001, q = 6.549). Increasing calcium beyond standard FSCV concentration (from 2.4 mM to 4.8 mM) did not significantly increase DA release (*p* > 0.05, q = 1.7). Peak height correlated across increasing calcium concentration for upward velocity (**Figure 2C**; n = 20, r^2^ = 0.7884, *p* < 0.0001, slope = 0.2857 ± 0.03489). This correlation was not as strong as that observed with baclofen and quinpirole experiments, possibly due to the use of multiple pulse stimulations which may increase variability due to recruitment of additional feedback mechanisms. Thus, from GPCR-mediated inhibition experiments, and calcium dependent release experiments, there is a clear relationship between release peak height and the upward velocity of release. This is not surprising since velocity simply and effectively quantifies the rising slope of an existing curve. Changes in upward velocity would only be expected if there were large increases in peak height associated with decreased dopamine clearance.

**Figure 2.**
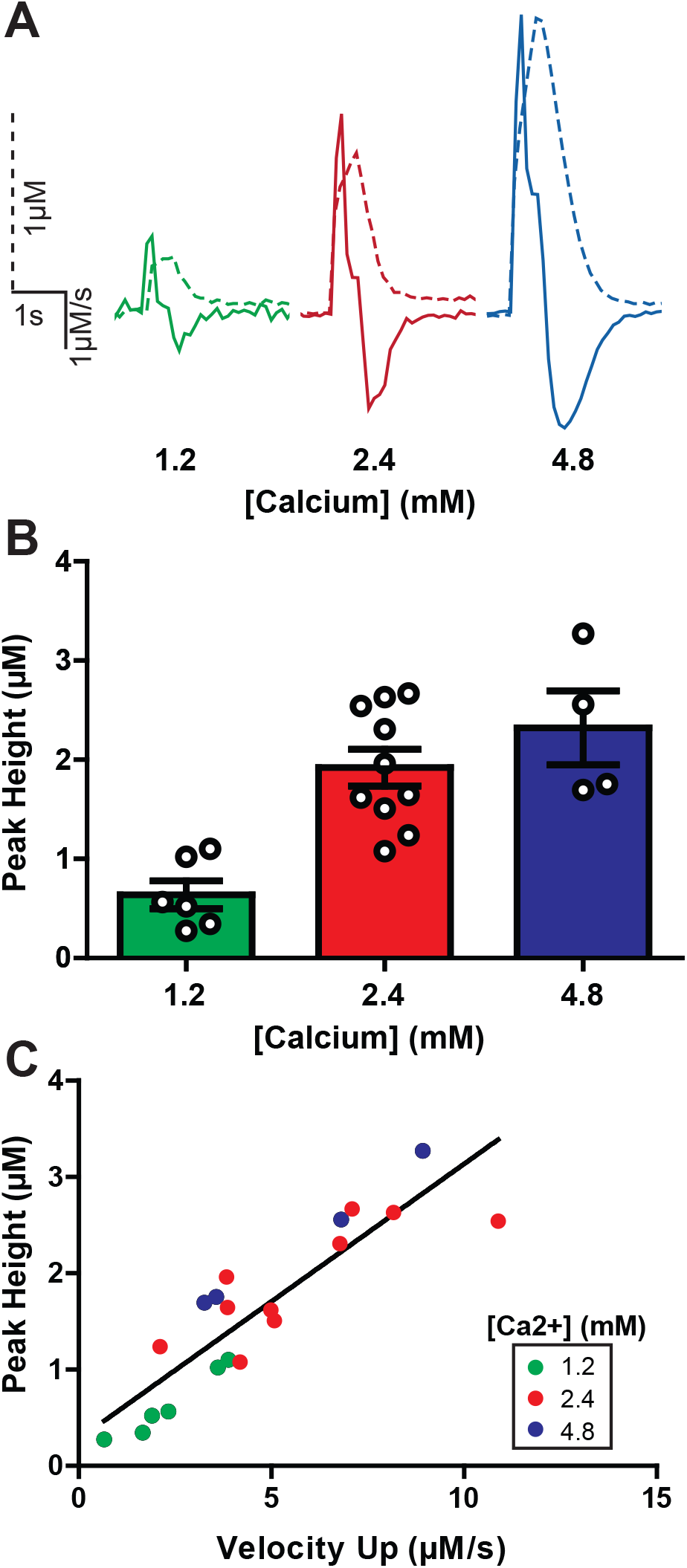
Upward velocity and peak height as measures of changes in DA release with calcium. A) A comparison of example dopamine traces (dotted lines) overlaid with the corresponding first derivative (solid lines) at increasing concentrations of calcium. All traces are scaled identically. B) Maximal evoked dopamine concentrations (μM) at increasing concentrations of calcium (mM). C) The ascending arm of the first derivative (Velocity Up) corresponds linearly with an increase in evoked dopamine release. Calcium concentrations for each measure are denoted by color.

### Downward velocity as a measure of dopamine transporter function

The predictive value of maximal downwards velocity was investigated in relation to the Michaelis-Menten value V_max_—an analytical value indicative of the maximal rate of dopamine clearance mediated by the DAT (Wightman 1988) (Yorgason *et al*. 2011a). The Michaelis-Menten V_max_ measure was obtained under baseline (non-drug) conditions in brain slices (*ex vivo*; **Figure 3A, B**) and in anesthetized mice (*in vivo*; **Figure 3C, D**). The traces of evoked dopamine release (1 pulse) and first derivatives with low (blue) and high (green) downward velocity values from *ex vivo* recordings are shown (**Figure 3A**). Maximal downward velocity and Michaelis-Menten V_max_ values in brain slices are correlated (**Figure 3B**; n = 8, r^2^ = 0.6239, slope = −0.1581 ± 0.05011). *In vivo* evoked (60 Hz, 30 pulses) dopamine release traces with low (orange) and high (pink) downward velocity rates are shown (**Figure 3C**). From *in vivo* experiments, downward velocity values correlated with Michaelis-Menten V_max_ values (**Figure 3D**; n = 9, r^2^ = 0.6699, slope = −0.8435 ± 0.2238). Interestingly, Michaelis-Menten V_max_ values in slice were much larger than *in vivo* values (*ex vivo*: 2.463 ± 0.472; *in vivo*: 1.025 ± 0.094 μM/s; Two tailed t-test: *t*_16_ = 2.987, *p* = 0.0087). In contrast, downward velocity values covered a similar range and were thus more comparable between *in vivo* and *ex vivo* conditions, though still significantly different (*ex vivo*: −1.05867 ± 0.09544; *in vivo*: −0.69785 ± 0.084399 μM/s; Two tailed t-test: *t*_14_ = −2.472, *p* = 0.02689). This suggests that the Michaelis-Menten modeling analysis has a bias towards faster values in slice compared to *in vivo* conditions, supporting the use of downward velocity for comparisons across recording conditions.

**Figure 3.**
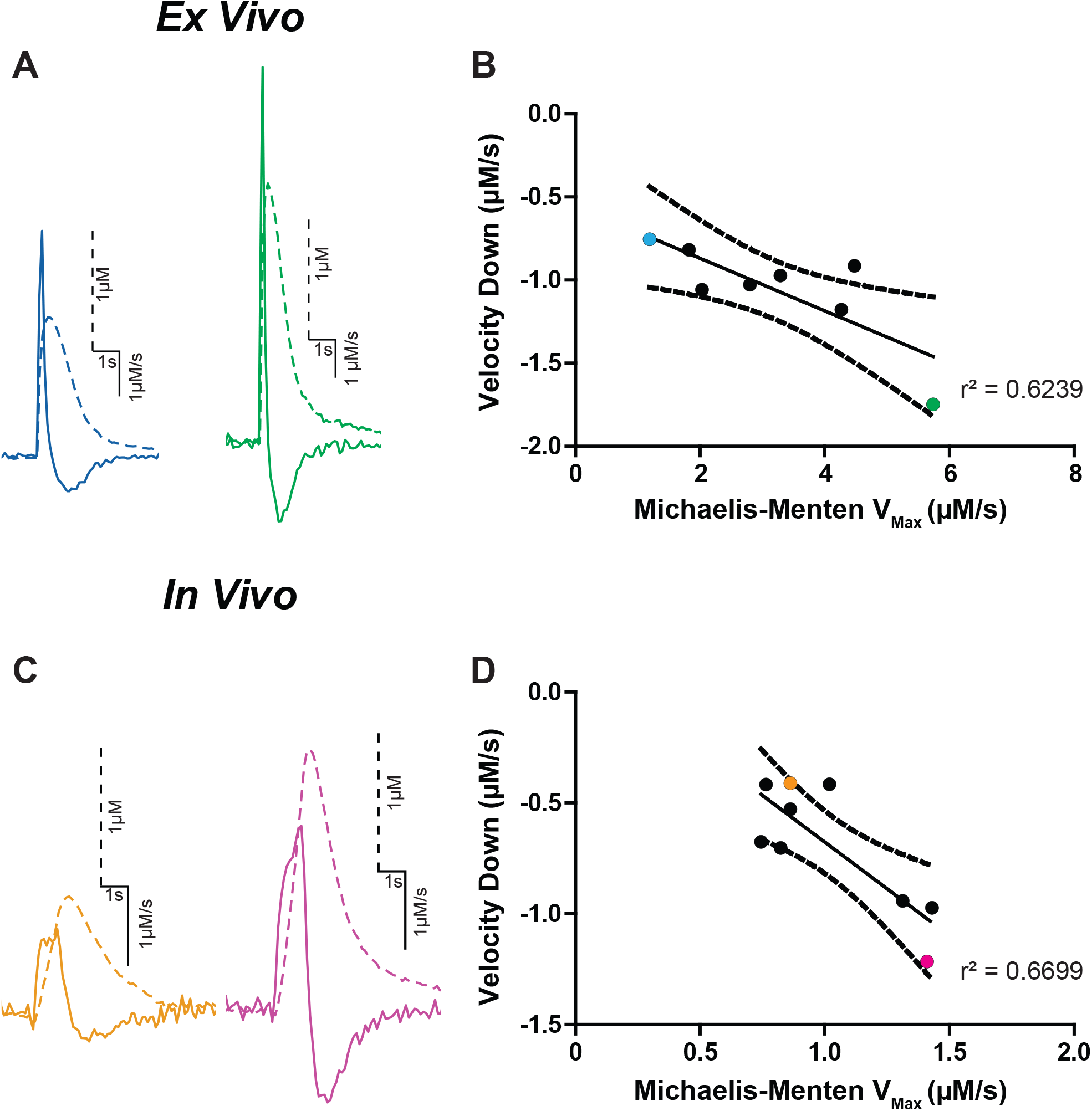
Downward velocity as a measure of dopamine transporter function. A) Evoked dopamine traces from *ex vivo* recordings. Traces are scaled individually. Dopamine concentration (dotted lines) and first derivative (solid line) are overlaid. Each trace is scaled individually. B) The V_max_ obtained using the Michaelis-Menten model is plotted against the descending arm of the first derivative (Velocity Down). Each dot represents one trace, with the colored dots corresponding to their respective colored traces in Figure 2A. C) Evoked dopamine traces from *in vivo* recordings. Traces are scaled individually. D) Michalis-Menten V_max_ compared to decreasing velocity from *in vivo* recordings. Colors correspond to their respective traces in Figure 2C.

### Downward velocity as a measure of dopamine clearance in DAT knockout mice

Downward velocity and release measures were examined in DATKO mice to determine how much the DAT contributes to each of these measures (**Figure 4**). Since downward velocity is an effective predictor at measuring DAT function in relation to the Michaelis-Menten Vmax measure, it was expected that this measure would be most affected by DAT removal. Furthermore, Michaelis-Menten parameters cannot be tested in DATKO mice due to violation of modeling parameters from complete lack of DAT enzymatic activity (Wightman & Zimmerman 1990). Dopamine signals were examined at three stimulation conditions which produce varying DA release concentrations (1 pulse, 5 pulses at 20Hz, 24 pulses at60Hz). Example dopamine release (**Figure 4A**) and velocity (**Figure 4B**) traces are shown. Upwards velocity positively correlated with peak height in WT mice (velocity: n = 46, r^2^ = 0.7027, *p* < 0.0001, slope = 1.92 ± 0.19) and in DATKO mice (velocity: n = 36, r^2^ = 0.6419,*p* < 0.0001, slope = 1.59 ± 0.2). The relationship between peak dopamine and upward velocity (across stimulation conditions) was examined, and no significant difference was observed between WT and DATKO mice (F_1,78_ = 1.38, *p* = 0.2431). Therefore, peak dopamine signals from multiple pulse stimulations relate to maximal upward velocity measures, and this relationship is maintained in the absence of the dopamine transporter. Two-way repeated measures ANOVA for peak release (**Figure 4C**) revealed a main effect of stimulation intensity (F_2,56_ = 186.74, *p* < 0.0000001), but no effect of mouse strain (F_1,56_ = 2.65, *p* = 0.115) and no significant interaction (F_2,56_ = 1.25, *p* = 0.294). For upward velocity (**Figure 4D-E**) there was a main effect of stimulation intensity (F_2,56_ = 68.48, *p* < 0.0000001), but no effect of mouse strain (F_1,56_ = 0.70, *p* = 0.408) and no significant interaction between these terms (F_2,56_ = 0.27, *p* = 0.766). Thus, detection of release as measured by peak and upward velocity is increased with higher stimulation intensities and is similar between WT and DATKO mice.

**Figure 4.**
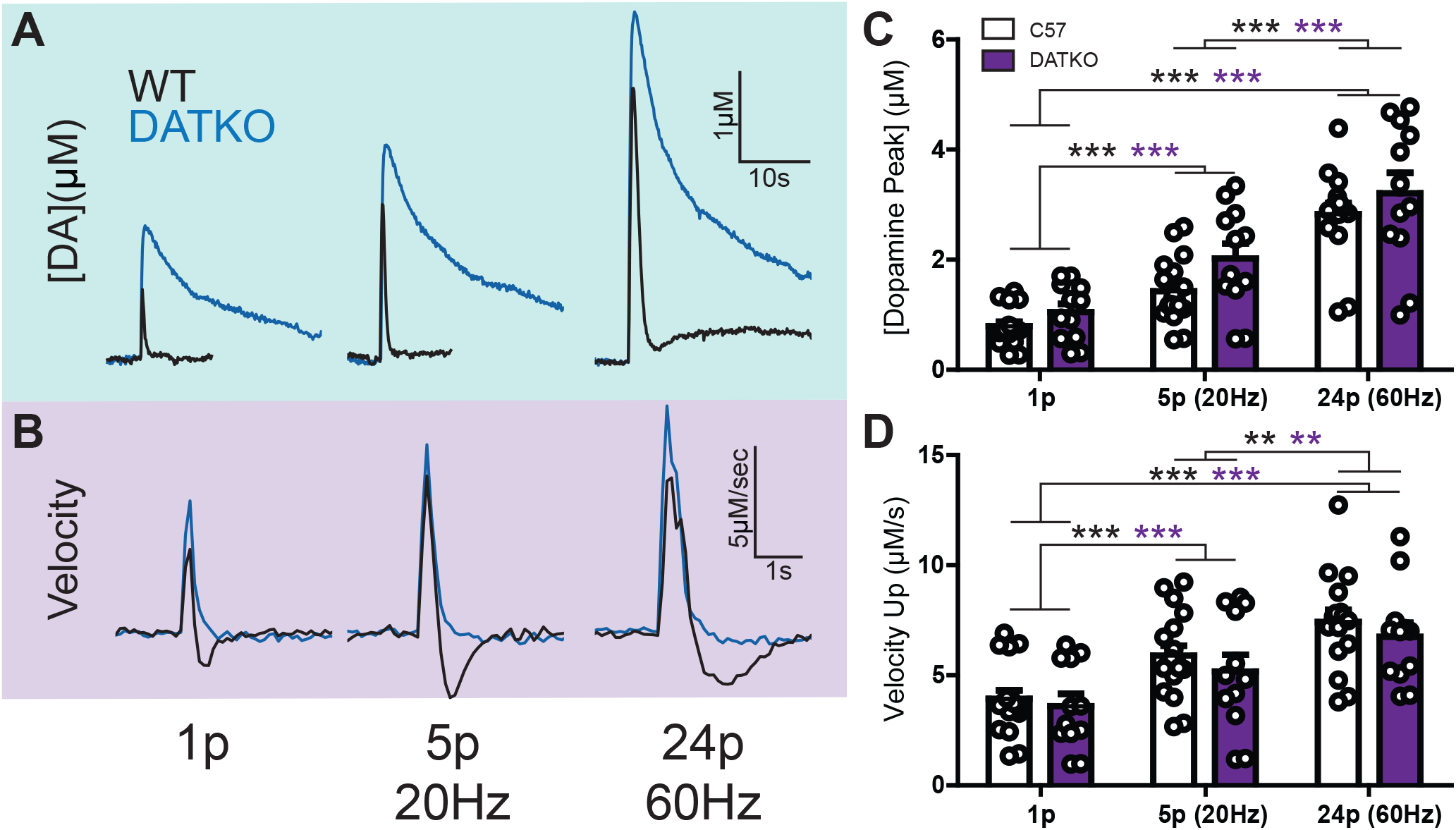
Upward velocity is not indicative of difference in uptake in DAT knockout mice. A) Evoked dopamine release concentrations compared between wild-type (black) and DAT knockout (blue) mice. Increasing pulse number and frequency induced higher release concentrations. Pulse protocols began with a single pulse (left), then increased to 5 pulses at 20 Hz (middle) and ended with 24 pulses at 60 Hz (right). B) The first derivative (velocity) of the dopamine traces in Figure 4A. C) Average peak dopamine concentrations for the three pulse conditions. Each dot represents a single pulse event. D) First derivative maximum (peak velocity) for the three pulse conditions. Each dot represents a single pulse event. [** represents *p* < 0.01; *** represents *p* < 0.001]

Dopamine clearance was evaluated using striatal slices from WT and DATKO mice (**Figure 5**). Average (±SEM) first derivative curves across an evoked release time course are shown (**Figure 5A**; 20 Hz, 5 pulse stimulation). There are clear differences in overlap for the downward velocity slope in these two strains. A subtraction between average dopamine velocity curves from WT and DATKO mice across stimulation conditions highlights the time course of velocity differences (**Figure 5B**). Interestingly, greater downward velocities were observed with higher stimulation intensity (compare green 5 pulse curve to black 1 pulse curve in **Figure 5B**), with greatest velocities observed in WT mice at the 60Hz 24 pulse stimulation conditions (WT 1p: −1.395 μM/s; WT 20Hz5p: −1.963 μM/s; WT 60Hz24p: −2.138 μM/s). Not surprisingly, longer duration stimulations also resulted in subtraction curves with longer duration downward velocities. These results show that DAT effects across time can be more pronounced with longer stimulations. Maximal downward velocity values across stimulation conditions and strains are shown (**Figure 5C**). Two-way repeated measures ANOVA for downwards velocity revealed a main effect of stimulation intensity (F_2,56_ = 23.27,*p* < 0.0000001) and mouse strain (F_1,56_ = 12.33, *p* = 0.0015), but no significant interaction between these terms (F_2,56_ = 1.58, *p* = 0.215). These DATKO experiments highlight the utility of downward velocity for measuring large changes in dopamine uptake by complete removal of the DAT.

**Figure 5.**
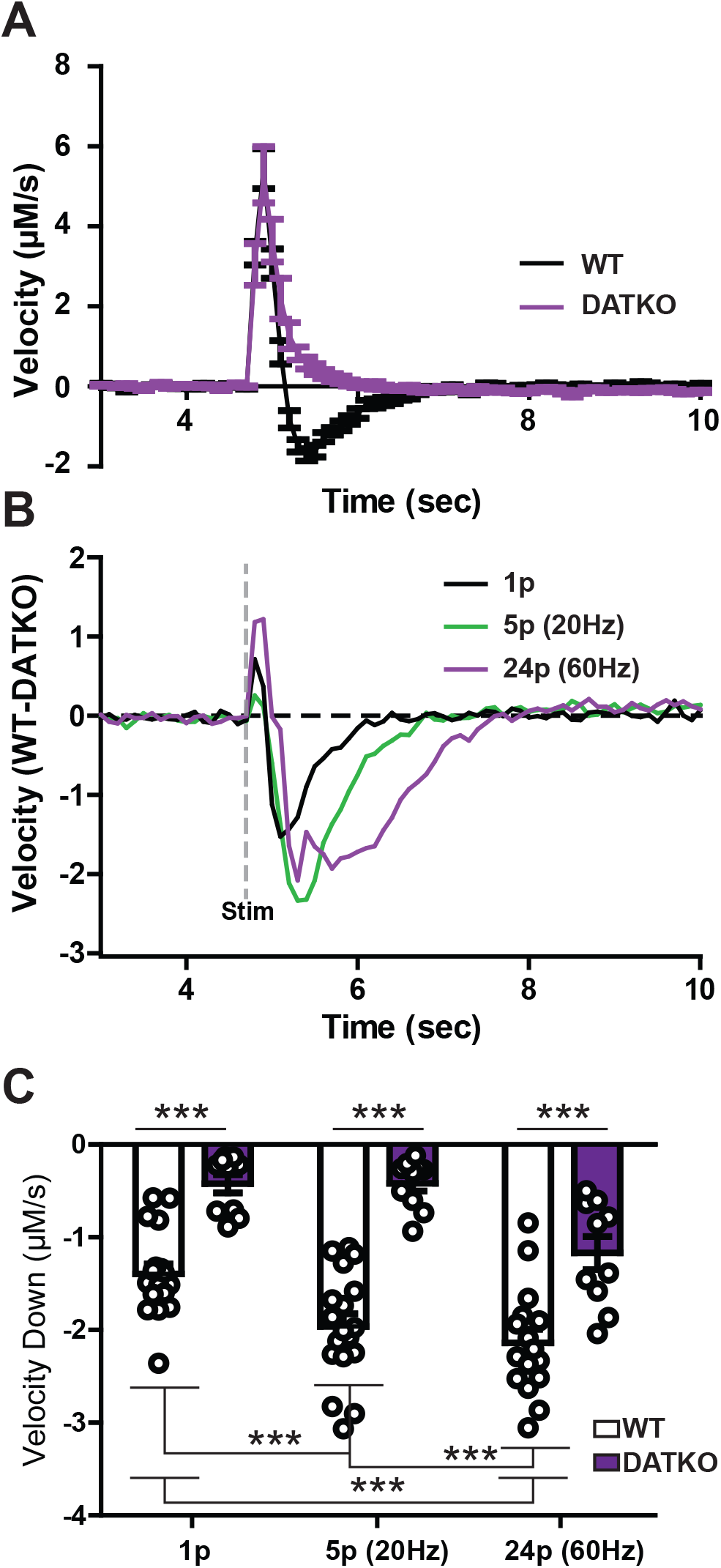
Downward velocity as a measure of dopamine clearance in DAT knockout mice. A) A comparison of wild-type (black line) and DAT knockout (purple line) velocities obtained via the first derivative of dopamine concentration release under the single pulse condition. Note the slow decrease in DATKO compared to the negative values in the WT mouse. B) A subtraction plot of a first derivative trace in a WT mouse minus a trace from the DATKO mouse for each pulse condition. An electrical stimulation was delivered at 5 seconds as denoted by the dashed grey line. C) Average minima for the first derivative (peak velocity down) compared by mouse type and pulse condition. Each dot denotes a single DA release event. [*** represents *p* < 0.001]

### Dopamine transporter blockers differential effects on release and uptake

The maximal downward velocity from the first derivative mathematically corresponds to the fastest time point where DA is being cleared, which is adjacent to the time point for the peak height (**Figure 1A**). Dopamine transporter inhibitors of the tropane family share a binding site with dopamine (Bisgaard *et al*. 2011) (Beuming *et al*. 2008) (Wang *et al*. 2015). Therefore, it is expected that due to competition effects, downward velocity will have varying sensitivity to DAT inhibitors that will be weakest at lowest concentrations and weakest with inhibitors with lower affinity. The effects of tropane DAT competitive blockers with varying affinities (WF23, PTT, Cocaine) on dopamine clearance was measured *ex vivo* (**Figure 6**). Representative release traces are shown for WF23 (**Figure 6A**), PTT (**Figure 6B**) and cocaine (**Figure 6C**). The highly potent WF23 (K_i_ = 0.1 nM) (Davies *et al*. 1994) elevated peak height (**Figure 6D**) across increasing concentrations (10 nM-3 μM), with greatest increases observed at the 300 nM concentration at ~458% of baseline. Similarly, PTT (K_i_ = 3 – 8.2 nM; 100 nM – 30 μM) (Bennett *et al*. 1995) (Letchworth *et al*. 2000) increased peak height with greatest effects at 1 μM (~282% of baseline). Cocaine (Ki = 120 – 189 nM; 300 nM-30 μM) (Gatley *et al*. 1996) (Katz *et al*. 2000) elevated peak height to a maximum of ~134% baseline at 1-3 μM. Uniquely, cocaine also decreased signals by ~49% from baseline at 30μM. Although WF23 and PTT both exhibited an inverted U-shaped curve, peak height never fell below baseline levels for these drugs. Two-way ANOVA on peak height data revealed a main effect of concentration (F_8,60_ = 6.28, *p* = 0.000007), but indicated no drug effect (F_2,60_ = 0.06, *p* = 0.942093 or interaction (F_1,60_=1.51, *p* = 0.125862).

**Figure 6.**
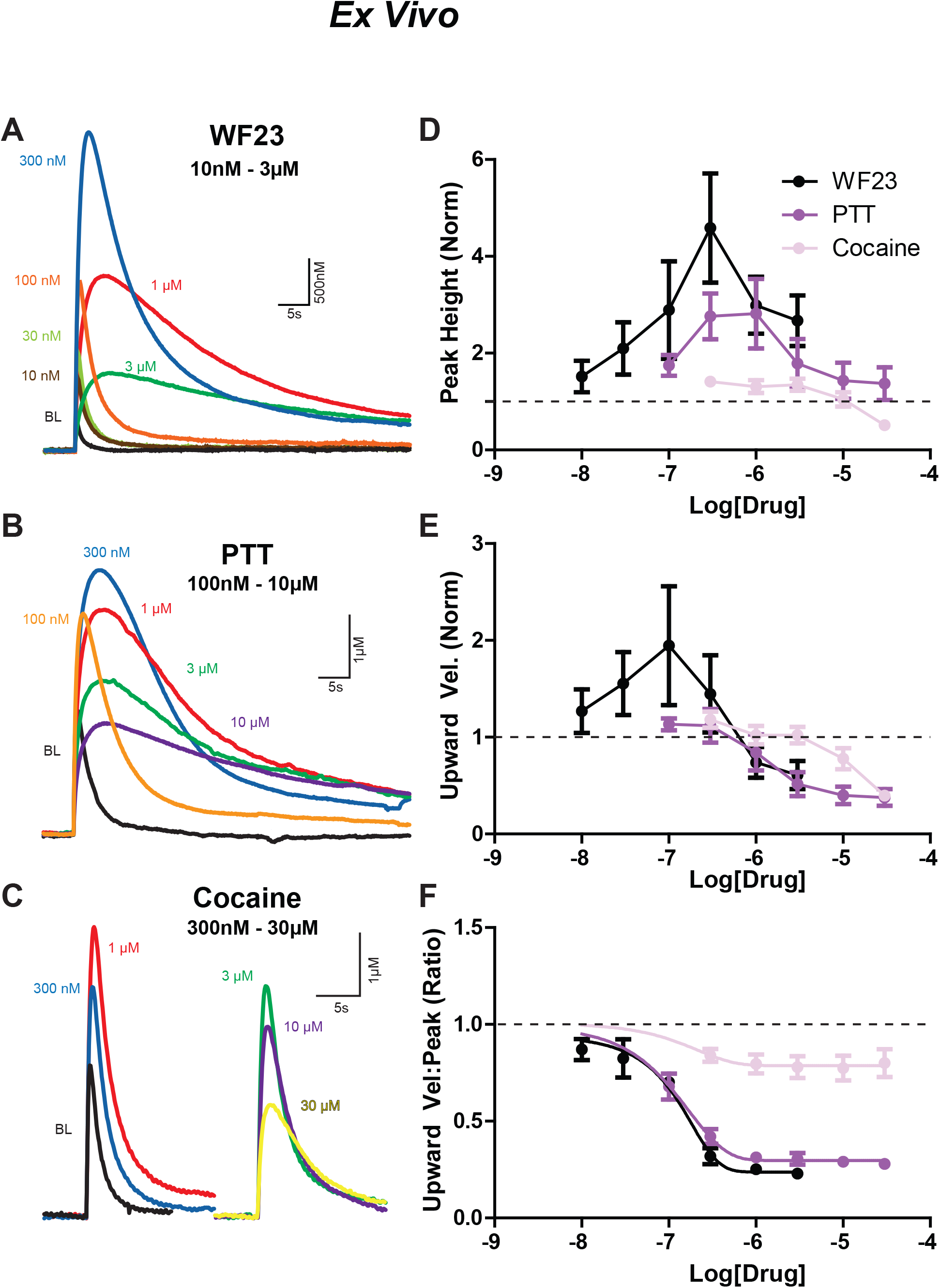
Effects of DAT blockers on DA release kinetics *ex vivo*. Traces from A) WF23 (10 nM – 3 μM), B) PTT (100 nM – 30 μM, and C) cocaine (300 nM – 30 μM) drug concentrations found in Figures 6D, E, and F, respectively. Signals denoted by color: baseline (black), 10 nM (brown), 30 nM (yellow-green), 100 nM (orange), 300 nM (blue), 1 μM (red), 3 μM (green), 30 μM (yellow). D) Comparison of peak height of signals across concentrations for WF23 (black), PTT (purple), and cocaine (pink). Similar comparisons across drug concentrations for E) upwards velocity and F) the ratio of upwards velocity to peak height. All measurements are normalized to baseline.

The upward velocity measures were affected differently than peak height by competitive DAT inhibitors. The greatest effects for upward velocity across the three drugs were at 100 nM for WF23 (~194%) and 300 nM concentrations for PTT (~112%) and cocaine (~118%; **Figure 6E**). Upward velocity reduced below baseline at >1 μM (WF23, PTT) and >10 μM (cocaine) concentrations. This is an important example where increases in peak height do not relate directly to upward velocity and suggests that the increase in peak height observed at the higher concentrations of inhibitors is likely due to a different mechanism, such as increased spread from distant dopamine terminals. This phenomenon can be observed in traces from WF23 and PTT experiments at high concentrations, where the peak height is larger than the baseline, and the time for that peak is delayed (rightward shifted), reflecting a slower upward velocity (**Figure 6A-B**). Two-way ANOVA on upward velocity data revealed a main effect of concentration (F_8,60_ = 4.19, *p* = 0.000497), no drug effect (F_2,60_ = 0.73, *p* = 0.499866), and no interaction (F_1,60_ =1.20, *p* = 0.291346).

Since the association between upward velocity and peak height is weaker in the presence of high concentrations of DAT blockers, this measure may be particularly useful in dissociating and interpreting mechanisms underlying increases in peak height that are not due to increased vesicular fusion. The ratio between upward velocity and peak height was examined across drug concentrations (**Figure 6F**) and exhibited a clear decrease for all DAT blockers, with greater efficacy and potency for WF23 and PTT over cocaine. Two-way ANOVA on the ratio between upward velocity and peak height data revealed a main effect of concentration (F_8,60_ = 70.35,*p* < 0.0000001), a main effect of drug (F_2,60_ = 12.16, *p* = 0.001054, and a significant interaction (F_1,60_ = 8.45, *p* < 0.0000001).

The effects of tropane analogs on downward velocity in striatal slices were examined next (**Figure 7**). First derivative traces are shown for WF23, PTT and cocaine (**Figure 7A-C**). All three drugs reduced downward velocity at high concentrations (**Figure 7D**), but effects were mixed at low (<100 nM) concentrations, with WF23 with some experiments exhibiting faster downward velocity (see example traces in **Figure 7A**). Therefore, WF23 effects were examined with (**Figure 7D**) and without (**inset**) experiments where velocity increased at low concentration ranges. For WF23 and PTT experiments, downward velocity was reduced for a similar range of concentrations (>100 nM), which was leftward shifted from cocaine, reflective of the increased potency of WF23 and PTT for the DAT over cocaine. Cocaine effects were modest (apparent effects at >1 μM), and did not plateau until ≥30 μM. These data suggest that downward velocity is an effective tool for measuring relative potency for competitive DAT inhibitors.

**Figure 7.**
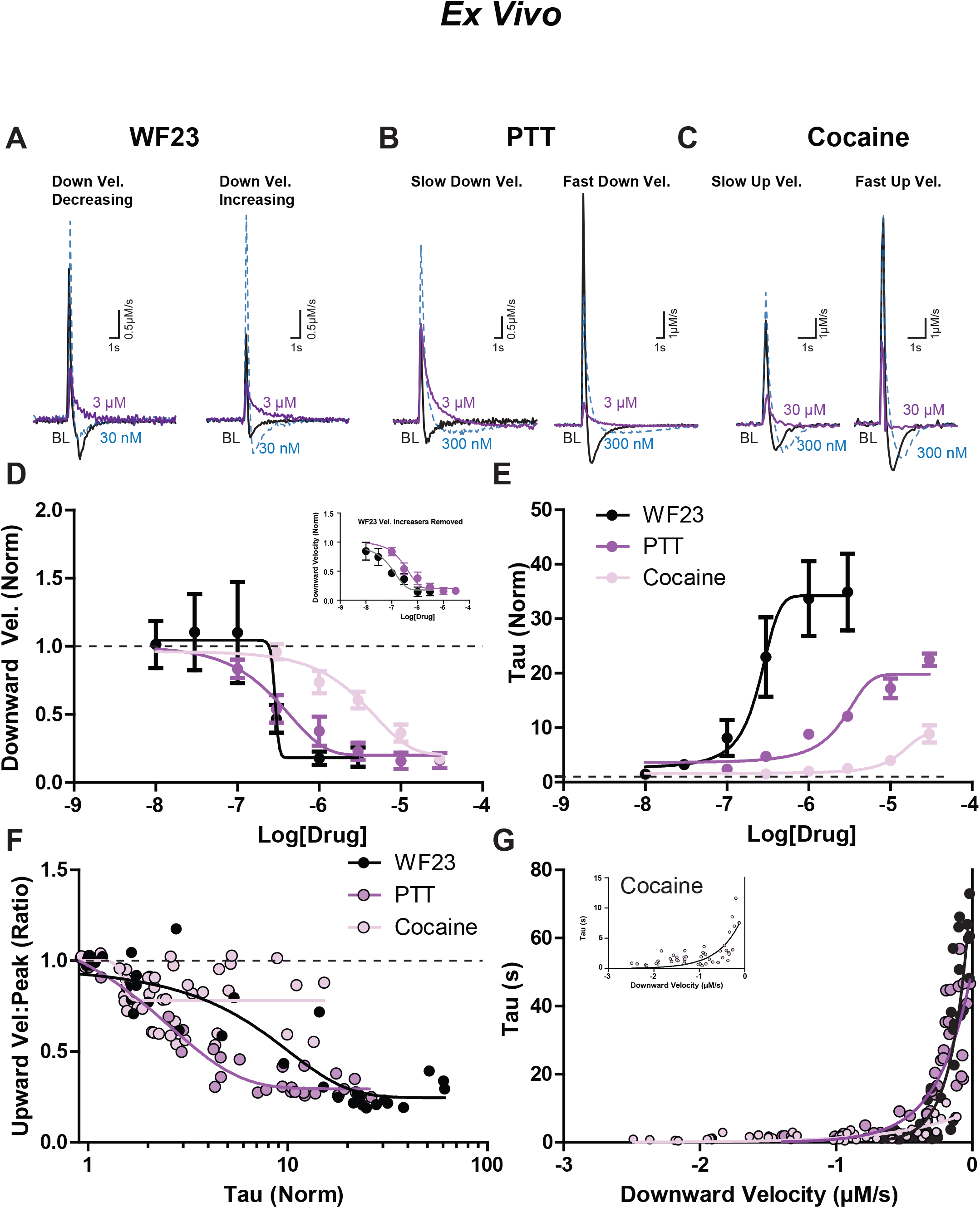
Effects of DAT blockers on DA uptake kinetics *ex vivo*. Example first derivative traces are shown under baseline conditions (black), 30 nM (blue), and 3uM (purple) for A) WF23, B) PTT, and C) cocaine. A) three experiments indicated a distinct decrease in downward velocity with addition of WF23 at a low concentration (left) while in two experiments the downward velocity increased (right). B) experiments with slower downward velocity at baseline (left) are shown next to signals with faster downward velocity at baseline (right). C) experiments with slower upward velocity at baseline (left) are depicted next to signals with fast upward velocity at baseline (right). D) Downward velocity (normalized) and E) tau (normalized) are compared across increasing concentrations of WF23 (black), PTT (purple), and cocaine (pink). F) Relationship between tau (normalized) and the ratio of upward velocity to peak height for each drug. G) Relationship between downward velocity (μM/s) and tau (sec) for each drug.

The Michaelis-Menten K_m_ is frequently used for describing effects of DAT inhibitors (Yorgason *et al*. 2016) (Calipari *et al*. 2014) (Siciliano *et al*. 2015) (Torres *et al*. 2021) and correlates strongly with the exponential decay measure tau (Yorgason *et al*. 2011a). Therefore, in order to establish the relationship between exponential decay and downward velocity, the effects of tropane analogs on tau were examined (**Figure 7E**). Tau was more sensitive to tropane analogs than downward velocity, with WF23 increasing tau with the greatest efficacy and potency (LogEC_50_ = 0.2418), followed by PTT (LogEC_50_ = 2.326) and cocaine (LogEC_50_ = 12.59). Two-way ANOVA on tau data revealed a main effect of concentration (F_8,60_ = 13.90,*p* < 0.0000001) and a significant interaction between these terms (F_16,60_ = 4.69,*p* <0.0001), but no main effect of drug (F_2,60_ = 2.51, *p* = 0.119840). Considering the increased sensitivity of tau over downward velocity, downward velocity appears more effective at measuring changes in dopamine uptake reflective of changes in transporter numbers (similar to V_max_), while tau appears more effective for detecting changes in apparent affinity of dopamine for the DAT (similar to K_m_).

Decreases in the upward velocity to peak height ratio (seen in **Figure 6F**) may be due to reduced dopamine clearance. Since tau is a sensitive measure of impaired clearance for competitive inhibition (**Figure 7E**), the relationship between these two values was examined. Upward velocity to peak ratio correlated to tau (**Figure 7F**) for WF23 (Spearman r = −0.8202, *p* < 0.0001) and PTT (Spearman r = −0.8619, *p* < 0.0001) but less for cocaine (Spearman r = −0.3802, *p* = 0.0077), likely reflecting cocaine’s lower efficacy at reducing the upward velocity to peak height ratio and lower efficacy on increasing tau. These data highlight the relationship between increases in DA signals and decreases in uptake which are most apparent as tau increases, but are different for each competitive inhibitor. The relationship between tau and downward velocity was further examined in the context of DAT inhibitors (**Figure 7G**). Tau values significantly correlated with downward velocity for WF23 (Spearman r = 0.7838,*p* < 0.0001) PTT (Spearman r = 0.8937, *p* < 0.0001) and cocaine (Spearman r = 0.6522, *p* < 0.0001), and tau was most strongly related at smaller downward velocities. Therefore, reductions in upward velocity to peak ratio are due in part to impaired clearance, and tau becomes more predictive of downward velocity in conditions where dopamine clearance is severely impaired by reuptake inhibitors.

The effects of potent competitive DAT inhibitors (WF23 and PTT) were tested *in vivo* in order to establish a measure sensitivity for these two different drugs. Experiments were performed across a 30-minute time course, and measures of peak height, upward and downward velocity, and tau were all examined (**Figure 8**). WF23 and PTT increased dopamine peak height for both drugs (**Figure 8A**; Two-way RM-ANOVA; Time effect: F_12,144_ = 8.015, *p* < 0.001; Drug effect: F_1,144_ = 1.616, *p* = 0.2278) with a significant interaction between drug and time variables (Interaction: F_12,144_ = 15.00, *p* = 0.0120). These drugs had a similar effect on upward velocity (**Figure 8A Inset**; Two-way RM-ANOVA; Time effect: F_12,144_ = 5.621, *p* < 0.0001; Drug effect: F_1,144_ = 1.152, *p* = 0.3042; Interaction: F_12,144_ = 1.945, *p* = 0.0336). The ratio between these measures decreased across time similarly for both drugs (**Figure 8B**; Two-way RM-ANOVA; Time effect: F_1,144_ = 3.004, *p* = 0.009; Drug effect: F_1,144_ = 2.485, *p* = 0.1409; Interaction: F_12,144_ = 3.012, *p* = 0.0009). These drugs also reduced clearance as measured by increases in tau, with clear differences between these drugs (**Figure 8C**; Two-way RM-ANOVA; Time effect: F_12,144_ = 8.015, *p* < 0.0001; Drug effect: F_1,144_ = 6.925, *p* = 0.0001; Interaction: F_12,144_ = 3.609, *p* = 0.0233). The upward velocity to peak ratio correlated with changes in tau (**Figure 8D**) for WF23 (Spearman r = −0.7282, *p* < 0.0001) and weakly with PTT (Spearman r = −0.2082, *p* = 0.0136). Interestingly, in some WF23 and PTT experiments the downward velocity increased initially (**Figure 8E**; during the first 20 minutes; Two-way RM-ANOVA; Time effect: F_12,144_ = 1.505, *p* = 0.1284; Drug effect: F_1,144_ = 0.07187, *p* = 0.7932; Interaction: F_12,144_ = 1.031, *p* = 0.4240), likely due to initial low concentration effects on increasing DAT activity which were also observed in slices (**Figure 7A, D**). When these increasing values are removed, there is a clear decrease in downward velocity induced by WF23 and PTT at the 5 second time points (**Figure 8E Inset**; Two-way RM-ANOVA; Time effect: F_12,144_ = 8.460,*p* < 0.0001; Drug effect: F_1,144_ = 0.7706, *p* = 0.4202; Interaction: F_12,144_ = 1.001, *p* = 0.4594). Similar to slice experiments, downward velocity from *in vivo* experiments correlated with tau (**Figure 8F**) for WF23 (Spearman r = 0.6884, *p* < 0.0001). However, this relationship was not maintained with PTT (Spearman r = 0.04761, *p* = 0.5764).

**Figure 8.**
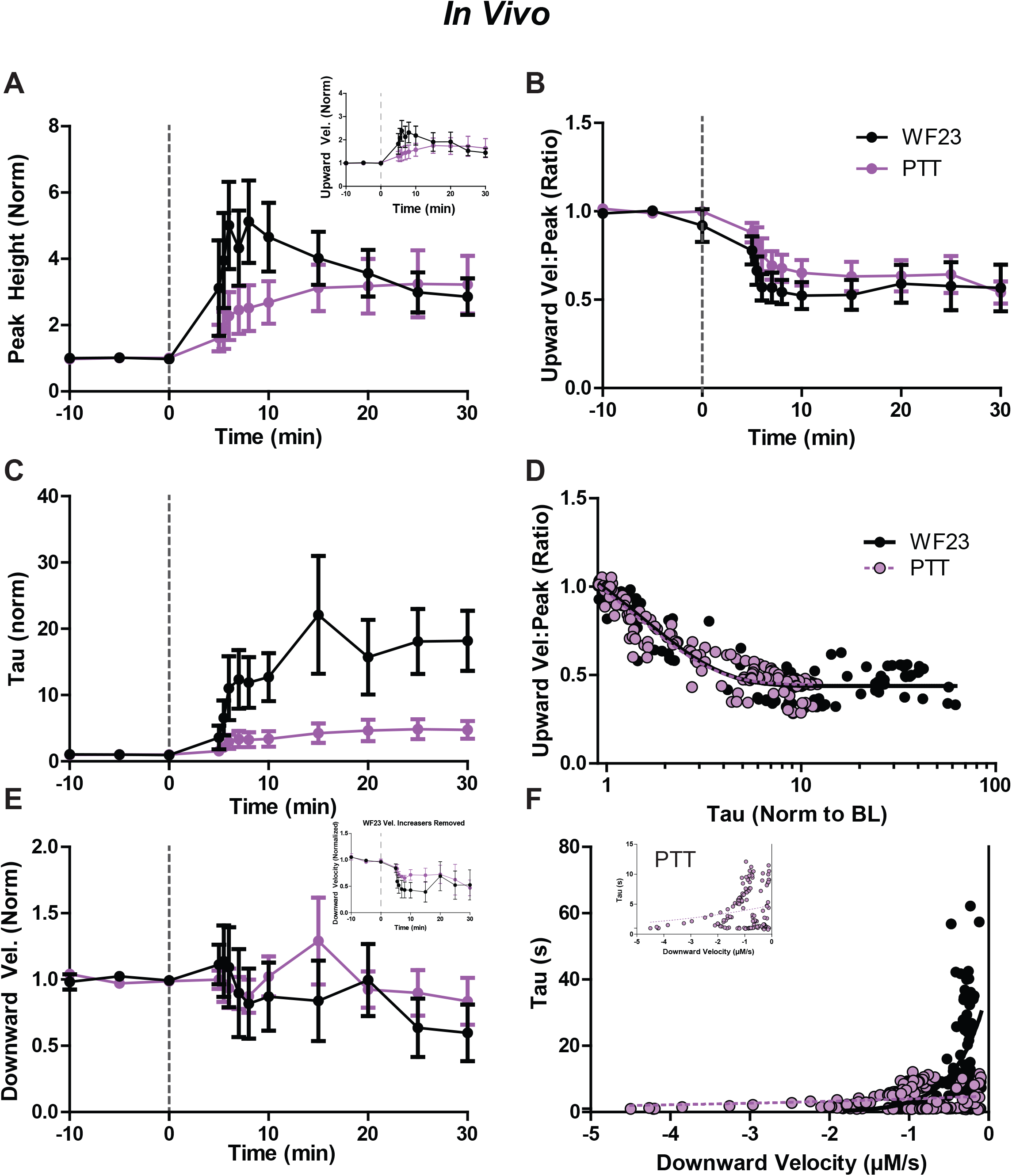
Effects of DAT blockers on DA release and uptake kinetics *in vivo*. Time course across 30 minutes (post-injection) of A) peak height, B) ratio of upward velocity to peak height, C) tau, and E) downward velocity of DA signals for WF23 (black) and PTT (purple). All values normalized to baseline. D) Comparison of tau (normalized) to ratio of upward velocity to peak height for each drug. F) Comparison of downward velocity (normalized) to tau (normalized) for each drug.

## Discussion

The present experiments examine measures of dopamine release and clearance. Dopamine release (determined by the peak height of signals) is reduced by application of GABA_B_ and D_2_ agonists and increased with enhanced calcium levels and greater stimulation intensity. This release measure correlated strongly with maximal upward velocity. Maximal downward velocity correlated with the Michaelis-Menten uptake measure of V_max_, an indirect measure of DAT expression, whereas the exponential decay measure tau related more to the Michaelis-Menten measure K_m_. Genetic and pharmacological blockade of the DAT resulted in concentration dependent reductions in the observed maximal downward velocity and increases in tau, with greatest sensitivity in the tau measure. The application of these measures to dopamine signals introduces an unbiased approach for measuring release and clearance, which is needed for improving analysis standardization across the field.

The relationship between drug induced changes in peak height and upwards velocity is complex and represents a concert between cellular mechanisms underlying release and clearance. In conditions where release is solely influenced, increases in peak height are tightly related to velocity increases. In contrast, for tropane concentration response experiments, peak height increased across drug concentrations, but the velocity of the rising peak varied greatly in a drug and concentration dependent manner (typically increasing at low and decreasing at high concentrations respectively). Peak height increases associated with slower upward velocities indicate the crude nature of peak height as a measure of release. The dissociation between these measures apparent in the presence of DAT blockers is indicative of the multitude of effects of these drugs on DA terminal function.

The modified Michaelis-Menten model used by voltammetrists calculates the amount of dopamine released ([DAp]) in the context of DAT enzymatic activity values (V_max_ and K_m_) (Wightman & Zimmerman 1990). This same model was previously used to measure cocaine’s DAT-blocking effects and comparisons were made across DA signal peak height, [DAp], K_m_ and the area under the curve (Yorgason *et al*. 2011a). From these previous experiments, peak height correlated strongly with [DAp] (r^2^=0.95) across cocaine concentrations (300 nM-30 μM). Herein, similar increases in peak height were observed across concentrations of DAT blockers, with increases above baseline for most concentrations, and slight reductions in dopamine peak with just cocaine at 30 μM (reduced by ~20%). As a comparison, upward velocity increased at concentrations ≤ 1 μM for WF23/PTT (peak increases of ~32-57% at 300 nM respectively) and ≤ 3 μM for cocaine (peak increases of ~33% at 300 nM). In contrast, upward velocity decreased at higher concentrations (~18% decreases with WF23 at 3 μM; ~33-42% decreases with PTT at 10-30 μM; ~13-48% decreases with cocaine at 10-30 μM). Together with our previous study, these data suggest that the [DAp] and DA peak height are not completely reflective of release, and interpretation is influenced in cases of reduced uptake where the peak height is increasing but upward velocity is not. In experiments where the potent DAT blockers PTT and WF23 were applied, peak dopamine release was increased over baseline values across concentrations, but with reduced upward velocity in the 3-30 μM range. This concentration range coincided with larger reductions in downward velocity. This suggests that the decrease in upwards velocity, but overall increase in peak, is due to increased detection of release from diffusion from distant compartments. Surprisingly, and in contrast to DAT blocker experiments, there were similar upward velocities observed between WTand DATKO mice. This finding was disconcerting because DATKO mice have been used heavily as a comparison group for studying clearance kinetics in the absence of the DAT (i.e. diffusion) (Jones *et al*. 1998) (Benoit-Marand *et al*. 2000). These results suggest that there are compensations in DATKO mice affecting release and clearance. Therefore, experiments examining DA diffusion may benefit from using these or other high affinity DAT blockers. These results also support an unknown adaptation in dopamine clearance—possibly secondary clearance mediated by the cholesterol sensitive organic cation transporter (Gasser 2019).

The diversity of observed effects of DAT inhibitors on peak height and velocity measures of release across concentrations are indicative of the multiple effects of DAT inhibitors on DA terminals. One of the mechanisms recruited during DAT blockade is activation of dopamine D2 type receptors (Kimmel *et al*. 2001). Dopamine D2-type receptors are inhibitory GPCRs, located on presynaptic dopamine terminals (as autoreceptors) and on postsynaptic neurons (as heteroreceptors) in the striatum. At the soma, D2 autoreceptors activate inhibitory effectors, including the G-protein-coupled inward rectifying potassium (GIRK) channels, to directly drive membrane hyperpolarization (Pillai *et al*. 1998). In vivo, cocaine increases dopamine levels (Greco & Garris 2003), resulting in D_2_ mediated inhibition of somatic activity (Einhorn *et al*. 1988). In a slice, evoked dopamine release is insensitive to D_2_ antagonists (Phillips *et al*. 2002) (Kennedy *et al*. 1992), demonstrating that extracellular tone is minimal in slice conditions. Fastscan controlled adsorption voltammetry (FSCAV) studies in slices have reported extracellular dopamine levels of ~11 nM in the striatum (Burrell *et al*. 2015) and ~40 nM in the SNc (Yee *et al*. 2019). Bath application of cocaine (10 μM) increased extracellular dopamine in the SNc by ~6.8 nM (Yee *et al*. 2019). There are currently no FSCAV reports on dopamine levels after cocaine in striatal slices. However, bath application of dopamine (1 μM) produces modest (~5- 10%) activation of D2 receptors (IC50 of 36 μM), and cocaine (10 μM) quadruples D2 sensitivity (IC50 at 9 μM) (Marcott *et al*. 2014). Several studies have shown D_2_ mediated inhibition of dopamine release with cocaine at high concentrations (10 μM) (Holroyd *et al*. 2015) (Adrover *et al*. 2014). We have observed D_2_ mediated inhibition of dopamine release by cocaine at concentrations as low as 1 μM (Yorgason *et al*. 2017). At low concentrations insufficient for inhibiting uptake (10 nM), cocaine has been shown to enhance D2 receptor stimulation possibly due to positive allosteric effects on D_2_ receptors (Ferraro *et al*. 2010) (Genedani *et al*. 2010) (Ferraro *et al*. 2012). Blocking D_2_ receptors in the presence of cocaine (1-10 μM) results in greater dopamine release (Yorgason *et al*. 2017) (Adrover *et al*. 2014) (Holroyd *et al*. 2015). Since dopamine release velocity reductions were observed herein in the presence of the D_2_ agonist quinpirole, and since DAT blockers examined are known to increase D_2_ activation, the decrease in velocity observed with DAT blockers can be attributed in part to D_2_ auto activation. In terminals, D_2_ activation will likely influence several effectors to attenuate dopamine release, including activation of Kv2.1 channels (Fulton *et al*. 2011), inhibition of snare proteins (Blackmer *et al*. 2001) and inhibition of synthesis proteins (i.e. tyrosine hydroxylase) (Chen *et al*. 2020). It is unknown if GABA_B_ receptors recruit this same machinery at dopamine terminals, or if some of cocaine’s effects are through enhancement of GABA_B_ effects, though it seems likely that adaptations from cocaine would affect GABA_B_ sensitivity considering the overlap in effector coupling for these receptors (Guatteo *et al*. 2004).Cholinergic interneurons (CINs) are a major regulator of dopamine activity, and electrically evoked dopamine release is initiated in part through release of acetylcholine and subsequent depolarization through activation of nicotinic acetylcholine receptors (nAChRs) (Yorgason *et al*. 2017) (Wang *et al*. 2014). Furthermore, nAChR allosteric activators and nAChR blockers enhance and reduce dopamine release respectively (Gao *et al*. 2019) (Yorgason *et al*. 2017). Also, the dopamine release process is dependent on sodium channel mediated depolarization on dopamine terminals (Yorgason *et al*. 2017), subsequent opening of voltage gated calcium channels (Brimblecombe *et al*. 2015) and calcium entry, which interacts with snare proteins to trigger vesicular fusion (Chen *et al*. 1999). Cocaine has been shown to block nAChR conductance at concentrations > 3 μM (Acevedo-Rodriguez *et al*. 2014). Mice lacking the β-2 nAChR subunit have reduced sensitivity to inhibition from cocaine (Acevedo-Rodriguez *et al*. 2014). In this latter study, peak reductions still occurred in β-2 knockout mice, but with a considerable reduction in sensitivity. Indeed, attenuated increases were observed in WT mice at concentrations > 2 μM concentrations, but in β −2 knockouts only at concentrations > 10 μM (Acevedo-Rodriguez *et al*. 2014). The inhibition observed at higher concentrations in β-2 knockout mice was attributed to cocaine’s well-known anesthetic effect (Acevedo-Rodriguez *et al*. 2014). Since WF23 and PTT share some structural homology with cocaine, it is likely that these drugs are acting through these several mechanisms to attenuate release. However, whether WF23 and PTT have nAChR or anesthetic effects at the tested concentrations remains unknown.

The downward velocity measure was highly indicative of DAT function. Downward velocity significantly correlated with Michaelis-Menten V_max_ values in vivo and in slice in WT mice and was considerably reduced in DATKO mice. Furthermore, downward velocity decreased after application of DAT blockers in slice and *in vivo*. Thus, downward velocity reliably predicts slower DAT kinetics in the presence of blockers. Since DATKO mice have no enzymatic function, the Michaelis-Menten V_max_ cannot be used in DATKO analysis. This highlights a utility of downward velocity in studying DA clearance without violating specific enzymatic conditions.

In addition to downward velocity, a measure of exponential decay (tau) was also applied to the data. Increasing concentrations of DAT blockers resulted in higher values of tau *ex vivo*. These higher values of tau corresponded with the relative potency of the DAT blocker (highest for WF23 and lowest for cocaine). Similarly, *in vivo* experiments revealed a large increase of tau for mice with WF23 injections and a much smaller increase for mice with PTT injections. This corresponds to the Michaelis-Menten K_m_, which increases with lower enzymatic affinity for a substrate (Wightman & Zimmerman 1990). Thus, the present study asserts the use of both downward velocity and tau as unbiased ways to analyze DAT kinetics independent of the Michaelis-Menten model.

